# Development of prokaryotic cell-free systems for synthetic biology

**DOI:** 10.1101/048710

**Authors:** Abel C. Chiao, Richard M. Murray, Zachary Z Sun

**Author notes:** Correspondence to: Zachary Z. Sun. NOTE: This is a technical report for future inclusion in work pending submission, review, and publication. Therefore, this work has not been peer-reviewed and is presented asis*.

## Abstract

Prokaryotic cell-free systems are currently heavily used for the production of protein that can be otherwise challenging to produce in cells. However, historically cell-free systems were used to explore natural phenomena before the advent of genetic modification and transformation technology. Recently, synthetic biology has seen a resurgence of this historical use of cell-free systems as a prototyping tool of synthetic and natural genetic circuits. For these cell-free systems to be effective prototyping tools, an understanding of cell-free system mechanics must be established that is not purely protein-expression driven. Here we discuss the development of *E. coli*-based cell-free systems, with an emphasis on documenting published extract and energy preparation methods into a uniform format. We also discuss additional considerations when applying cell-free systems to synthetic biology.

## INTRODUCTION

Cell-free systems have historically been a fundamental tool for biological research. This was partially a necessity, as transformation technologies to introduce DNA into *E. coli* (Mandel & Higa, 1970) and modern recombinant DNA technology (Smith & Welcox, 1970) were not available until 1970. As a result, many scientific phenomena were limited to probing in crude lysates. The most-well known is Nirenberg’s use of a cell-free system to decipher the genetic code for the Nobel Prize in Medicine in 1968 (Nirenberg & Matthaei, 1961).

When recombinant DNA technology became mainstream and working in cellular hosts the norm, cell-free systems were relegated to the production of hard-to-produce or high-value proteins such as antibodies (Ryabova, Desplancqh, & Spirin, 1997; Yin et al., 2012) or cytotoxic agents (Martemyanov, Shirokov, Kurnasov, Gudkov, & Spirin, 2001; Salehi et al., 2016). Significant field focus has been on producing large amounts of proteins, either by engineering of the cell-free system itself (Kigawa, Yabuki, Yoshida, Tsutsui, & Ito, 1999) or through assisted methods of production (Spirin, Baranov, Ryabova, & Ovodov, 1988a). These methods all utilize the ability of cell-free systems to efficiently produce protein without interference from cellular growth and metabolism. In addition, many systems are driven by T7 RNA polymerase expression (Krieg & Melton, 1987) to encourage as much protein production as possible. Completely “synthetic” cell-free systems from purified components (Shimizu et al., 2001) have also been developed for hard-to-produce proteins.

There has been a recent resurgence of using cell-free systems as a fundamental tool. An overview contrasting this approach to utilizing systems for expression is given in **Figure 1**. The goal is similar to original applications probing biological phenomena, but motivated by modern-day synthetic biology tools of DNA sequencing, synthesis, and assembly. The first implementation of this was in 2003, with genetic circuits in cell-free (Noireaux, Bar-Ziv, & Libchaber, 2003), followed by the high expression of native sigma70 promoters (Shin & Noireaux, 2010) and the implementation of a panel of native circuits (Shin & Noireaux, 2012). By uncoupling protein expression activity from cell-growth requirements and opening the system to external manipulation and perturbation, cell-free is increasingly being used as a “prototyping environment,” or an environment for testing hypotheses before final implementation (in a cell, or a cell-free environment) (Niederholtmeyer, Sun, Hori, & Yeung, 2015; Takahashi et al., 2014). The emphasis in this application is less on protein production and more on the data collected from the cell-free system itself.

**Figure 1.**
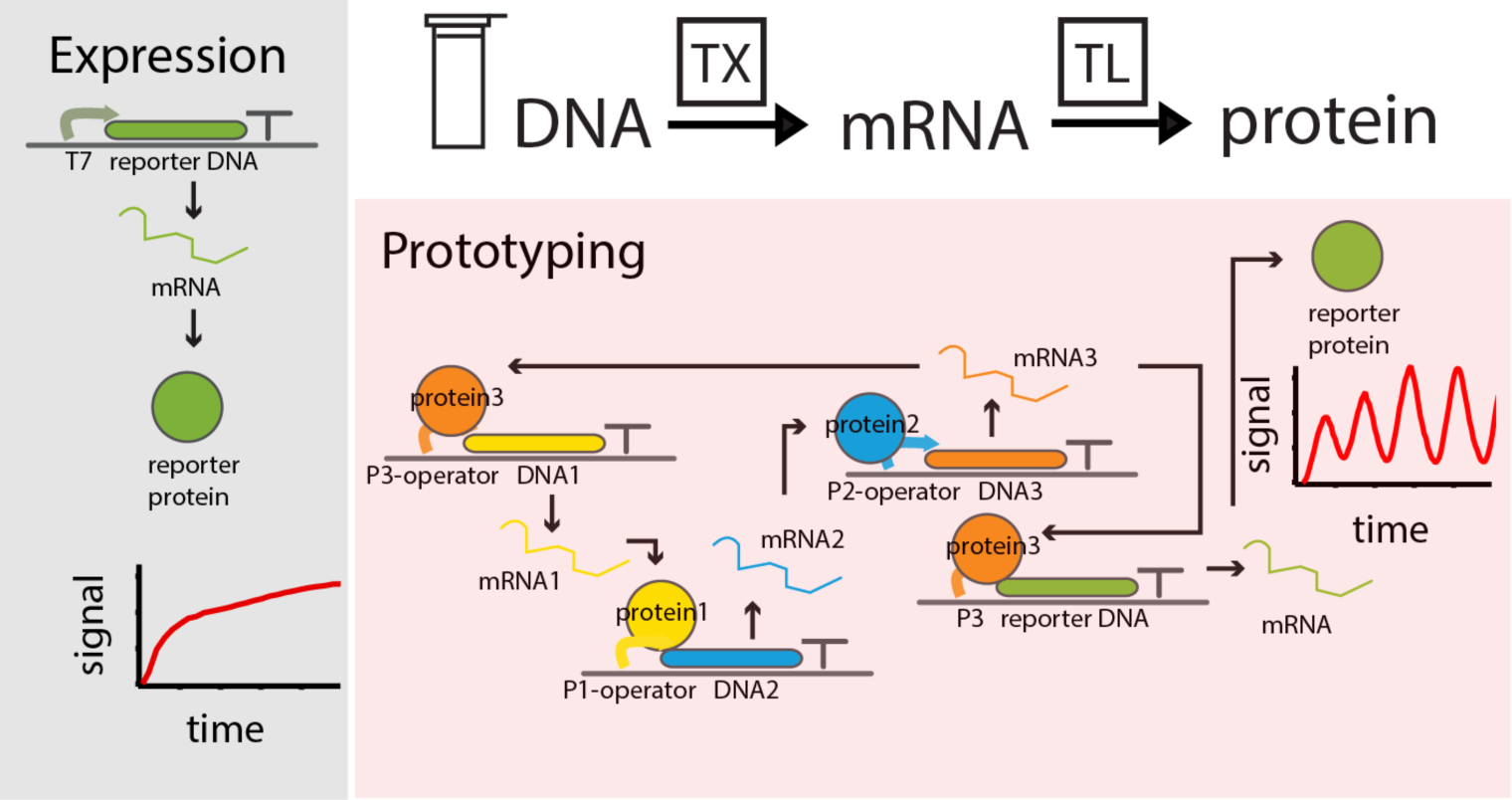
Overview of cell-free expression process. Execution is split into expression (left) and prototyping (right). On the right, the prototyping of a 3-node oscillator is represented.

In this review, we focus on the development and application of cell-free systems in synthetic biology. This diverges from, but builds off of previous in-depth reviews that take a broad-focus of cell-free systems as an expression platform (Carlson, Gan, Hodgman, & Jewett, 2012; Hodgman & Jewett, 2012; James R Swartz, 2012; Jim Swartz, 2006) or focus on engineering in cell-free (Takahashi et al., 2015). In doing so, we will explore the extensive prior research in cell-free system production and energy regeneration, as well as methods of executing cell-free reactions.

## *E. coli* EXTRACT PREPARATION METHODS

Since the earliest method for the cell-free synthesis of proteins was published by Nirenberg in 1961 (Nirenberg & Matthaei, 1961), numerous modified and improved protocols have been created. A detailed system published by Zubay in 1973 became what is considered to be the standard “S30” protocol upon which subsequent protocols were based (Spirin & Swartz, 2008; Zubay, 1973).

Generally, these protocols share a common schematic with the following features: a specific strain of *E. coli* upon which the extract is based, a strategy for cell culture, an elected method for cellular lysis, a centrifugation step to clarify the lysate, a heat incubation (run-off), dialysis, and post-dialysis clarification. Although this remains a fairly faithful general representation of the extract preparation process, none of these parameters were left unexplored in later iterations of the cell-free system and accordingly, current protocols do deviate from this paradigm. We parse extract production into the component pieces specified and evaluate the developments targeting each step (**Figure 2**).

**Figure 2.**
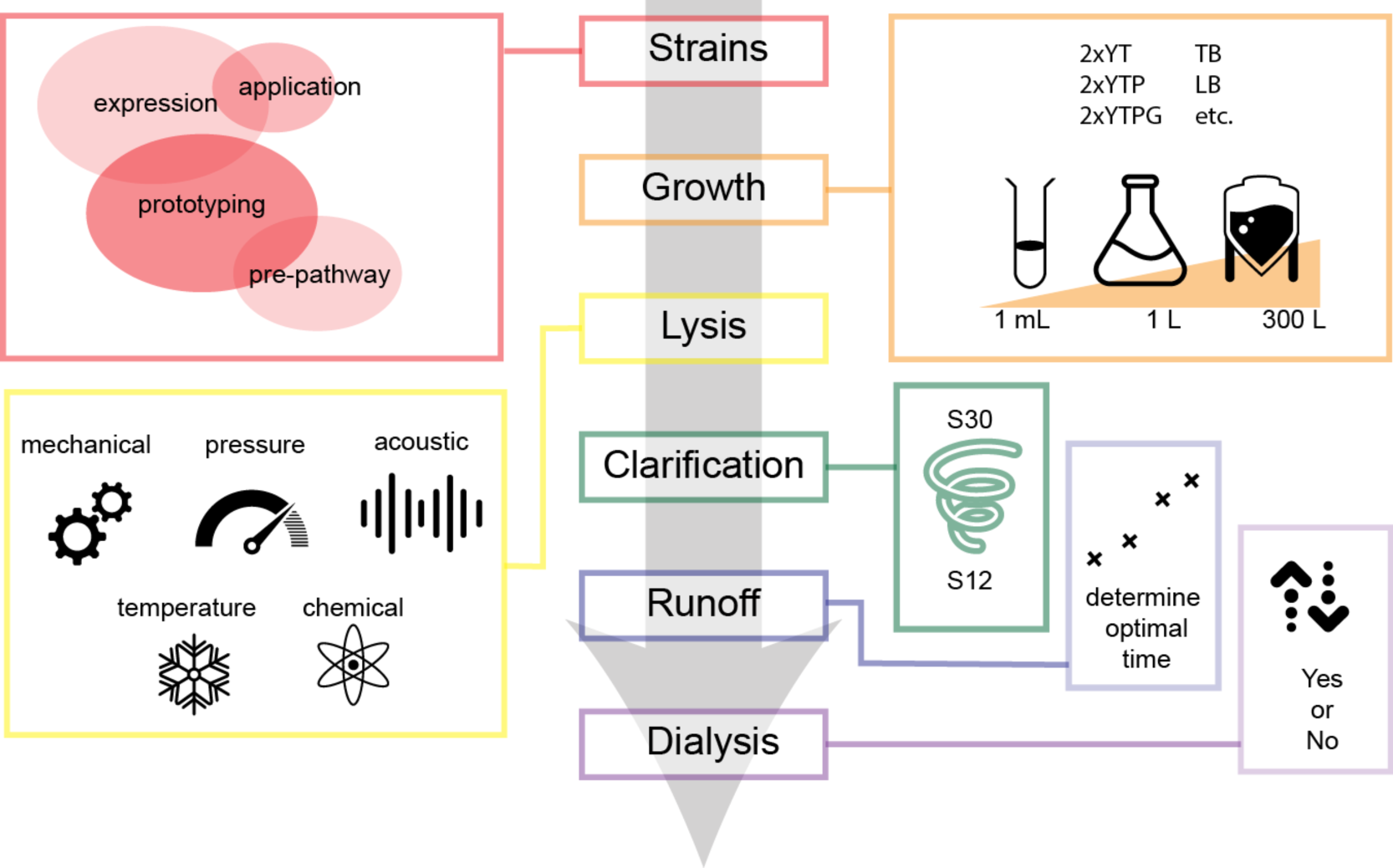
Flow-chart schematic for extract preparation. Preparation is divided into: strain (red), growth conditions (orange), lysis (yellow), clarification (green), runoff determination (blue), and dialysis (purple).

### Strain selection

Selection of model organism is critical as different *E. coli* strains offer a variety of different advantages over one another. Strains are differentiated by their engineered traits, some of which are very conducive to the controlled expression of protein. Choice of organism presents the first unit of modularity in the construction of cell-free systems. Engineered properties critical to the development of new applications can be directly conferred from engineered strains to their extracts. *E. coli* extracts have been successfully produced from a number of different cell strains. We divide strain selection into 4 overlapping areas: protein production-optimized strains, application-specific strains, generic strains, and pre-programmed pathway strains. Common genes overexpressed and under-expressed are in **Table 1**, and common strains are in **Table 2**.

**Table 1.** Genes commonly over-expressed or under-expressed in engineered cell-free strains. Citations indicate where more information about the gene in the context of cell-free can be found.

| Gene | Description | Citation |
| --- | --- | --- |
| **ackA+** | acetate kinase, added to increase yield | (Airen, 2011) |
| **csdA-** | cold shock degradosome protein, removed to prevent mRNA decay during preparation | (Hong et al., 2015) |
| **dnaJ+** | chaperone protein, added to assist folding with dnaK, grpE | (Kang et al., 2005) |
| **dnaK+** | chaperone protein, added to assist folding with dnaJ, grpE | (Kang et al., 2005) |
| **dsbC+** | disulfide isomerase, added for disulfide bond formation | (J. Yang, Kanter, Voloshin, Levy, & Swartz, 2004; Yin & Swartz, 2004) |
| **ef-tu+** | translation factor, added to increase yields (most abundant protein in cell, potentially rate-limiting) | (Airen, 2011; Woodrow & Swartz, 2007) |
| **endA-** | endonuclease, removed for plasmid stability | (Michel-Reydellet et al., 2004) |
| **gamS+** | lambda gam, added to protect linear DNA | (Michel-Reydellet et al., 2005; Sitaraman et al., 2004; Sun et al., 2014) |
| **gorB-** | glutathione reductase, removed to prevent disulfide bond persistence | (Kang et al., 2005; Knapp et al., 2007) |
| **groEJ+** | chaperone protein, added to assist folding with groEL+ | (Kang et al., 2005) |
| **groEL+** | chaperone protein, added to assist folding with groEJ+ | (Kang et al., 2005) |
| **grpE+** | heat shock protein, added to assist folding with dnaJ, dnaK | (Kang et al., 2005) |
| **gshA-** | glutamate-cysteine ligase, removed to stabilize cysteine | (Calhoun & Swartz, 2006) |
| **hchA+** | chaperone protein, added to increase solubility and yield | (Airen, 2011) |
| **ibpA+** | small heat shock protein (chaperone), added to increase solubility and yield | (Airen, 2011) |
| **ibpB+** | small heat shock protein (chaperone), added to increase solubility and yield | (Airen, 2011) |
| **lf-1+** | initiation factor 1, added to increase yield | (Airen, 2011) |
| **lf-2+** | initiation factor 2, added to increase yield | (Airen, 2011) |
| **lf-3+** | initiation factor 3, added to increase yield | (Airen, 2011; Kim A Woodrow et al., 2006; Woodrow & Swartz, 2007) US 20130316397 |
| **lacI-** | lacI repressor, removed to prevent interference with lacI-expressing circuits | (Niederholtmeyer et al., 2015) |
| **mazF-** | mazF toxin, removed to prevent mRNA degradation at 'ACA' sites | (Hong et al., 2015) |
| **met+** | P1 selection marker, engineering scar | (Michel-Reydellet et al., 2004) |
| **recD-** | recD, removed to protect linear DNA (ineffective) | (Michel-Reydellet et al., 2005) |
| **rna-** | RNAse A, removed for RNA stability | (Gesteland, 1966; D.-M. Kim et al., 1996) |
| **rnb-** | RNAse II, removed for RNA stability | (Hong et al., 2015; Woodrow & Swartz, 2007) |
| **rpfA-** | release factor 1, removed to encourage nsAA incorporation | (Hong et al., 2014) |
| **sdaA-** | serine deaminase, removed to stabilize serine | (Michel-Reydellet et al., 2004) |
| **sdaB-** | serine deaminase, removed to stabilize serine | (Michel-Reydellet et al., 2004) |
| **speA-** | arginine decarboxylase, removed to stabilize arginine | (Michel-Reydellet et al., 2004) |
| **tnaA-** | tryptophanase, removed to stabilize tryptophan | (Michel-Reydellet et al., 2004) |
| **tonA-** | outer membrane protein, engineering scar | (Michel-Reydellet et al., 2004) |
| **trxB-** | thioredoxin reductase, removed post-growth with HA tag to | (Kang et al., 2005; Knapp et al., |
| HA | prevent disulfide bond persistence | 2007) |

**Table 2.** Commonly used strains, with genotypes. Citations indicate originally developed locations and/or application

| Strain | Genotype | Citation |
| --- | --- | --- |
| **BL21-Rosetta2** | F<sup>-</sup> *ompT hsdS<sub>B</sub>(r<sub>B</sub><sup>-</sup> m<sub>B</sub><sup>-</sup>) gal dcm* (DE3) pRARE2 (Novagen) | (Shin & Noireaux, 2010; 2012; Sun et al., 2014) |
| **JS006** | MG1655 *araC-* *lacI-* | (Niederholtmeyer et al., 2015; Stricker et al., 2008) |
| **K12-A19** | *rna-* *gdhA2* *relA1* *spoT* *metB1* | (Gesteland, 1966; D.-M. Kim et al., 1996) |
| **KC1** | A19 *speA-* *tnaA-* *tonA-* *endA-* *sdaA-* *sdaB-* *met+* | (Michel-Reydellet et al., 2004) |
| **KC6** | KC1 *gshA-* | (Calhoun & Swartz, 2005a) |
| **KG6-der.** | KC6 *rnb-* *ackA+* *ef-tu+* *hchA+* *ibpA+* *ibpB+* *if-1+* *if-2+* *if-3+* | US20130316397 |
| **KGK10** | KC6 *gorB-* *trxB-HA* | (Knapp et al., 2007; Knapp & Swartz, 2007; Yin et al., 2012; Zawada et al., 2011) |
| **NMR1** | A19 *endA-* *met+* | (Michel-Reydellet et al., 2004) |
| **NMR2** | A19 *speA-* *tnaA-* *tonA-* *endA-* *met+* | (Michel-Reydellet et al., 2004) |
| **NMR4** | A19 *recD-* *endA-* *met+* | (Michel-Reydellet et al., 2005) |
| **NMR5** | A19 *lambda-phage<>recBCD* *met+* | (Michel-Reydellet et al., 2005) |
| **S30BL/Dna** | BL21(DE3) *dnaK/J+* *grpE+* | (Kang et al., 2005) |
| **S30BL/DsbC** | BL21(DE3) *dsbC+* | (Kang et al., 2005) |
| **S30BL/GroE** | BL21(DE3) *groEL/ES+* | (Kang et al., 2005) |
| **S30OB** | F<sup>-</sup> *omp ThsdS<sub>B</sub>(r<sub>B</sub><sup>-</sup> m<sub>B</sub><sup>-</sup>) gal dcm lacY1 ahpC* (DE3) *gor522:: Tn10 trxB* (Novagen) | (Kang et al., 2005) |
| **S30OB/Dna** | S30OB *dnaK/J+* *grpE+* | (Kang et al., 2005) |
| **S30OB/DsbC** | S30OB *dsbC+* | (Kang et al., 2005) |
| **S30OB/GroE** | S30OB *groEL/ES+* | (Kang et al., 2005) |

#### Protein production optimized strains

Increasing protein production yield has been one of the main focuses of cell-free optimization. Many of the strain modification findings in this area were pioneered by Swartz and colleagues from 2004 onwards. Earlier efforts on engineering cell-free systems focused on utilizing physical systems (D. M. Kim & Choi, 1996; Spirin, Baranov, Ryabova, Ovodov, & Alakhov, 1988b)) and engineering energy regeneration (Kigawa et al., 1999; Ryabova, Vinokurov, Shekhovtsova, Alakhov, & Spirin, 1995). However, in 2004 a seminal paper from Swartz et al. described deletions in genes encoding for amino acid degradation enzymes, thereby stabilizing amino acid supply and protein production (Michel-Reydellet, Calhoun, & Swartz, 2004). The paper identified four limiting amino acids: arginine, serine, tryptophan, and cysteine. Arginine was stabilized by removing *speA*, a gene encoding for a arginine decarboxylase, thereby inhibiting the conversion of arginine to putrescine. Serine was stabilized by removing serine deaminases *sdaA* and *sdaB*, inhibiting the conversion of serine to pyruvate. Tryptophan was stabilized by removing *tnaA.* There was an additional attempt to stabilize cysteine, but deletions in *tnaA* and *yfhQ* failed to achieve desired results. A follow-up paper identified a deletion in gshA, a glutamate-cysteine ligase, as the cysteine degradation culprit (Calhoun & Swartz, 2006). The resulting strain was named KC6 (Calhoun & Swartz, 2005a; Michel-Reydellet et al., 2004).

To stabilize templates off of which DNA can be translated, the lambda-phage cluster has also been inserted into strains made into cell-free lysates (Michel-Reydellet, Woodrow, & Swartz, 2005), creating the NMR5 strain. The insertion of lambda-phase cluster represented one of the first strain-engineering attempts at stabilizing linear DNA. Although the cluster was identified as *exo* and *beta* in (Michel-Reydellet et al., 2005), earlier efforts revealed *gam*, when added in purified form, to be the main RecBCD inhibitor and linear DNA stabilizer (Sitaraman et al., 2004).

Separately, work from the Kim (HJ) lab in 2005 identified strains with the overexpression of molecular chaperones capable of reducing aggregation and improve solubility of eukaryotic proteins such as human erythropoietin (Kang et al., 2005). The work inserted plasmids to overexpress chaperone and heat-shock genes *groEL/ES, dnaK/J* and *grpE*, or *dsbC.* Interestingly, the Kim group also explored the creation of extracts from the Origami strain (Novagen) that encourages disulfide bond formation. The roles of proteins *trxB, gor*, and *dsbC* would for later formally explored in the context of disulfide bond formation in (Knapp, Goerke, & Swartz, 2007).

With the success in engineering amino acid stability, high-throughput approaches for determining positive and negative factors to cell-free expression was explored. In a first attempt, Woodrow et al. expressed 55 genes from *E. coli* off of linear DNA templates in NMR5, and demonstrated gene expression (Kim A Woodrow, Isoken O Airen, & Swartz, 2006). This work was followed by an expression of 49 genes affecting transcription, folding, energy, and cell-division, coupled to a selective degradation of linear templates with DpnII (on methylation pattern) and a subsequent analysis of cell-free yields (Woodrow & Swartz, 2007). In a final iteration, Airen (in unpublished but peer-reviewed thesis work) expressed 3,789 *E. coli* open reading frames, identifying 79 positive effectors and 60 negative effectors (Airen, 2011). Using this information on negative effectors, 4 mutant strains were made that, when combined with (1) supplementation with positive effectors, (2) stabilization of pH, (3) substrate replenishment, and (4) mRNA stabilization were able to increase expression 3-4-fold. While strains with 4 negative effectors removed, *pnp, rnb, raiA*, and *mazG*, did not result in significant increased expression, supplementation in *ibpA, ibpB, if-1, if-2, if-3*, and *ef-tu* demonstrated increased yields (Airen, 2011; Woodrow & Swartz, 2007). Negative effectors *rna, rnb, csdA, mazF*, and *endA* have also been removed from newer MAGE-recoded (H. H. Wang et al., 2009) strains, resulting in increased fluorescent protein yield. *rna* and *rnb* code for RNAses, csdA for a cold-shock protein that degrades mRNA, *mazF* for a RNA-degrading toxin, and *endA* for a dsDNA endonuclease (Hong et al., 2015).

#### Application specific strains

Strain modifications have also been explored to enable the expression of proteins with disulfide bonds. Disulfide bonds are a common feature of mammalian proteins, but are difficult to implement in cell-free due to rapid reduction *in vitro* (Jim Swartz, 2006). While iodoacetamide treatment can inactive thiols responsible for reducing disulfide bonds (Yin & Swartz, 2004), the treatment globally targets ‐SH groups and can result in non-specific inactivation of critical enzymes (such as DsbC and G-3PDH) (Knapp et al., 2007). A workaround was through the creation of a deletion mutant of *trxB* (thioredoxin reductase) and *gor* (glutathione reductase) and supplementation with DsbC. Critically, *trxB* is tagged with a hemagglutinin tag to allow for it to be present during cell growth but removed after cell-free processing, as a double *trxB gor* knockout causes *ahpC* to mutate to a potent disulfide reductase (Knapp et al., 2007). It is noted that this genotype closely represents the Origami strain (Novagen) that contains knockouts of *trxB* and *gor* with suppressor mutations in *ahpC*, and was demonstrated successfully for cell-free production two years prior (Kang et al., 2005). The resulting strain (KGK10) or findings from engineering the strain form the basis for current production efforts of disulfide bond proteins. Commercially, Sutro Biopharma utilizes variants of the strain for producing cytokine rhGM-CSF at 200L scale (Zawada, Yin, Steiner, & Yang, 2011) and producing antibody fragment light and heavy chains (Yin et al., 2012).

#### Generic strains

While strains can be specifically engineered for protein production or for specific applications, there is also a focus on using “generic” strains for cell-free prototyping. Reasons for using generic strains rather than specialized strains include: (1) Lack of need of high protein expression, and (2) the desire to maintain prototyping fidelity between *in vitro* and *in vivo.* Generic strains such as K19, first introduced in 1966 (Gesteland, 1966), have been commonly used (Kigawa et al., 1999; D.-M. Kim, Kigawa, Choi, & Yokoyama, 1996), as well as MRE-600 (Spirin et al., 1988a), BL21-derivatives (CP strains (Kigawa, Yabuki, Matsuda, Matsuda, Nakajima, Tanaka, & Yokoyama, 2004a), Rosetta strains ((de los Santos, Meyerowitz, Mayo, & Murray, 2015; Shin & Noireaux, 2012; Sitaraman et al., 2004; Sun et al., 2013; Sun, Yeung, Hayes, Noireaux, & Murray, 2014; Takahashi et al., 2014)), DE3 strains ((Karim & Jewett, 2016; Kwon & Jewett, 2015; Kwon et al., 2013), Origami strains ((Kang et al., 2005)), and K12 MG1655 (Kwon & Jewett, 2015). These generic strains can be chosen for favorable properties of growth; for example, the Rosetta derivatives provide rare tRNAs, DE3-derivatives provide T7 RNA Polymerase, BL21-derivatives are optimized for protein production, and Origami derivatives optimize for disulfide bond formation with *txrB* and *gor* deletions. However, selection does not need to be limited to widely recognized substrains. For example, cells with lacI, araC, and tetR knockouts such as JS006 (Stricker et al., 2008) have been made into extracts to build oscillators that require exogenous lacI (Niederholtmeyer et al., 2015). In the reverse case, ExpressIQ (lacIQ) has been used to shut-down operons that are lacI sensitive (Sun, Kim, Singhal, & Murray, 2015). Commercially, cells optimized for 1,4-BDO production were used by Genomatica as the starting strain for lysis, to test hypotheses of expression efficiency (Fischer, 2016; Schilling, 2015). If using cell-free as a prototyping platform, where the data collected from cell-free systems is critical, the selection of strain is driven by the final *in vivo* implementation.

#### Pre-programmed pathway strains

Recently, strains with complete pathways already present have also been made into cell-free systems for the purposes of driving production of a specific product. This is distinct from directed cell-free synthesis of products by the combination of separate cell-free lysates, each with a single enzyme to drive catalysis, as exemplified by Zhang (YHP) and colleagues ((Rollin et al., 2015; Y. Wang, Huang, Sathitsuksanoh, Zhu, & Zhang, 2011).

Greenlight Biosciences has pioneered a unique method to produce cell-free systems pre-programmed with metabolic pathways of the product of interest, where energy flux is solely directed towards producing the product (and not towards cellular growth). This is achieved by compartmentalizing the cell into a cytoplasm and a periplasm, where the cytoplasm contains the pathway of interest without a key enzyme and the periplasm contains the key enzyme and proteases against tagged proteins that are essential for cellular growth and function (but divert metabolic flux *in vitro)* (James R Swartz, 2012). Upon lysis, both compartments are brought together. The protease can then degrade the tagged growth-related proteins, while the key enzyme can run the pathway. This process can be achieved by engineering the strain to have protein degradation tags on growth-related proteins and periplasm-export tags on key enzymes.

### Growth conditions

#### Growth volume

Historically, fermenters have been used to produce cell biomass. The original protocols utilized fermenters of up to 10 L in size to grow cells (Zubay, 1973). Building off of this, Swartz et al. demonstrated a 10 L scale (20 g/L wet pellet cell mass), which produced similar cell-free protein yield as shake-flask growth (Zawada & Swartz, 2005), but with the advantage of denser OD collection. The same protocol is cited by Sutro Biopharma in (Zawada et al., 2011), but utilizing a 200 L bioreactor also custom-retrofitted with baffles. In both cases, feed rates of glucose are controlled to prevent acetate accumulation. Fermenters can be used to scale up biomass production, but suffer from increased labor and monitoring needed to collect data.

In lieu, growth can be conducted on a shake-flask scale (1 L of cell culture in a 2.8 L - 4 L Erlenmeyer flask), which yields about 1-2 mL of crude extract per L (Sun et al., 2013). Shake-flasks allow for quick production of biomass without fermenter maintenance. Protocols are focused on maintaining fast growth and aeration before capture at culture mid-log phase, and thus typically use baffled flasks. Examples of protocols using shake-flasks can be found at (Kigawa, Yabuki, Matsuda, Matsuda, Nakajima, Tanaka, & Yokoyama, 2004b; Sun et al., 2013; W. C. Yang, Patel, Wong, & Swartz, 2012).

For smaller volumes, a recent protocol by Kwon and Jewett demonstrates the first rapid production of cell-free at the 10 mL culture tube, allowing for the rapid exploration of ~100 strains per day using basic, readily available equipment (sonicator, small shaker, tabletop centrifuge) (Kwon & Jewett, 2015). Expression levels from the 10 mL scale produce comparable protein to the 10 L scale. By allowing for small-scale but high throughput production, Kwon and Jewett’s protocol scales cell-free expression for exploring multiple rapidly-engineered strains or conditions.

#### Growth media

While older protocols utilized 28°C growth (Zubay, 1973), the current standard protocol utilizes 37°C growth to encourage rapid protein production. This is driven by the knowledge that ribosome concentration correlates directly with growth rate (Bosdriesz, Molenaar, Teusink, & Bruggeman, 2015). There is evidence that temperature, effecting growth rate, has a direct correlation with extract productivity. In particular, Kigawa and colleagues have found that extracts have a linear productivity from 20°C to 37°C of growth, where CAT production yield at 20°C is 66% that of 37°C (Seki, Matsuda, Yokoyama, & Kigawa, 2008). Similarly, Nakano’s lab identified amino-acid supplemented growth conditions that enabled cell-free growth of A19 at 42°C, yielding 40% more CAT production yield compared to 37°C growth (Yamane, Ikeda, Nagasaka, & Nakano, 2005). However, to date we are not aware of other protocols utilizing 42°C growth.

Growth medias vary from extract to extract perpetration, although in general medias used are complex, non-defined mixtures such as LB, 2xYT, etc. (Spirin & Swartz, 2008). Medias can be supplemented for limiting reagents; for example, asparagine, glutamine, and tryptophan (Yamane et al., 2005) is added to a complex media to encourage faster growth. For specific applications such as fermenter growth, glucose and amino acid concentration can be selectively monitored and fed to prevent acetate accumulation (Zawada & Swartz, 2005). In addition, in 2000 Kim and Choi identified the addition of phosphate and glucose to a 2xYT media (named 2xYT-PG) to be suppressive of phosphatase activity in the resulting extracts (R. G. Kim & Choi, 2000). Phosphatase activity was found to consume energy sources PEP and amino acid cysteine. The group’s working hypothesis was supplementation of phosphate and glucose would prevent the cell from making its own phosphatases to produce inorganic phosphate. This media forms the basis for most modern cell-free preparations. It is noted that cell-free formulations for prototyping to-date have removed glucose from the media (Caschera & Noireaux, 2015; Shin & Noireaux, 2010; Sun et al., 2013) for unknown reasons.

### Lysis methods

Following culture, a number of options exist for lysing the cells and harvesting. Considerations of note when selecting a lysis method are lysis efficiency, scaling potential, ease of use, and preservation of native cell components. Different lysis techniques have different pros and cons and consequently, different protocols have cell disruption methods tailored to their applications. There has been relatively little innovation in this area; reviews addressing general cell lysis, such as one from 1986, provide good overviews of the different methods (Chisti & Moo-Young, 1986). We divide our discussion of lysis methods into five categories: mechanical (non-pressure), mechanical (pressure), acoustic, temperature, and chemical.

#### Mechanical (non-pressure-based) lysis

Mechanical methods utilize a grinding mechanism of action. These techniques involve the agitation of a suspension of cells in the presence of ceramic/glass beads, the motion of which results in crushing and grinding forces that break apart the cells and efficiently shears DNA (Miller, Bryant, Madsen, & Ghiorse, 1999). Industrial scale bead-mills have been deployed for cellular lysis (Chisti & Moo-Young, 1986), although the use of “Bead-beater” type desktop devices have been preferred (Thompson & Chassy, 1981) and adopted in cell-free protocols (Kigawa, Yabuki, Matsuda, Matsuda, Nakajima, Tanaka, & Yokoyama, 2004b; Shrestha, Holland, & Bundy, 2012; Sun et al., 2013). Beads are easily separated from the lysate by centrifugation/filtering and no expensive equipment is required, greatly reducing the financial barrier of entry into cell-free biology. The protocol also has utility in lysing non-E. *coli* such as cyanobacteria (Mehta, Evitt, & Swartz, 2015) and environmental samples from soil (Yeates, Gillings, Davison, Altavilla, & Veal, 1998). To maintain high protein concentrations necessary for cell-free expression, beads can also be filtered out of solutions post-processing (Sun et al., 2013). As is the case with all mechanical lysis methods, localized sample heating that may denature native proteins is a concern in bead beating methods (Shrestha et al., 2012). This problem is circumvented by limiting lysis to short bursts and by incubating the samples on ice between bursts (Sun et al., 2013). Perhaps the largest drawback to this method of lysis is difficulty of scaling up to larger volumes as this method is typically conducted in 1-2 mL tubes. Bead mills present one avenue for scale-up but requires manual loading of extract. The ability to work in small volumes, however, is conducive to producing multiple distinct, small batches. Bead-beating is useful for studies requiring the production of multiple different extract batches, as is the case when conducting protocol modifications.

#### Mechanical (pressure-based) lysis

High pressure disruption mechanisms such as impinge homogenizers are among the earliest and most widely utilized methods for lysing cells for the purpose of preparing extracts (Chisti & Moo-Young, 1986). These work by forcing cell suspensions through a narrow aperture under high pressure. The high-velocity flow of cells either impinges on an opposite high-pressure stream of cells or a rigid valve/nozzle surface. The resulting shear rates and rapid decompression are thought to be critically important in the formation of inverted membrane vesicles in the resultant extracts (Jewett, Calhoun, Voloshin, Wuu, & Swartz, 2008). Because the enzymes essential to the oxidative phosphorylation pathway must be membrane-associated to function, the possibility of their presence is worth considering when selecting a lysis method. Access to the oxidative phosphorylation pathway potentially increases the metabolic efficiency of extracts, enabling more economical and productive strategies for powering the transcription-translation machinery.

For *E. coli* extracts, different types of impinge homogenizers are currently in use, ranging from French Press-style homogenization (Caschera & Noireaux, 2014; D. M. Kim & Choi, 1996; T. W. Kim et al., 2006) to Avestin™-type homogenization (Jewett et al., 2008; Liu, Zawada, & Swartz, 2005; Sitaraman et al., 2004; W. C. Yang et al., 2012). Both types of homogenizers allow for scaling of batches; French-press homogenizers scale up to the size of the press (typically 30 mL), while Avestin©-type homogenizers allow for feeding of cell biomass.

#### Acoustic lysis

Sonication, or acoustic lysis, relies on ultrasound energy (15-20kHz) to disrupt cells in solution. The mechanism of lysis is thought to be related to cavitation, a phenomena where microbubbles form at nucleation sites, absorb energy and burst, releasing mechanical shock waves that disrupts the cell wall and can shear DNA (Chisti & Moo-Young, 1986). There are relatively few examples of sonication being used as a lysis method for *E. coli* cell-free protein synthesis, with an early example failing due to “sample heating and difficulty of management” (Kigawa, Yabuki, Matsuda, Matsuda, Nakajima, Tanaka, & Yokoyama, 2004b). In 2012, Bundy and colleagues re-attempted sonication as a lysis technique, and were able to successfully demonstrate protein yields comparable to that of high-pressure homogenization, albeit with significant optimization of the sonication burst times and cooling times (Shrestha et al., 2012). In this study, temperature was also not shown to be a damaging factor. This was followed by a study from Kwon and Jewett optimizing energy input to cell-strain and processing volume, which found a surprising strain-dependence (Kwon & Jewett, 2015). It is anticipated that sonication will be studied further for *E. coli* cell-free systems. Like beadbeating, benefits of sonication include low startup costs and the ability to work with very small volumes.

#### Temperature based-lysis

Temperature-based lysis relies on freeze-thaw cycles to disrupt cellular membranes, and is one of the easiest methods of cellular disruption for producing purified proteins (Johnson & Hecht, 1994; Ron, Kohler, & Davis, 1966). This lysis can take place with or without enzymatic or chemical assistance such as lysosome. If successful, the method does not require advanced materials (other than liquid nitrogen or −80 C storage). However, freeze-thaw has not been demonstrated successfully for *E. coli* cell-free systems, with no appreciable expression detected despite a 99.6%-99.9% lysis efficiency (Shrestha et al., 2012). This is relatively surprising, as freeze-thaw in 20% glycerol has been demonstrated for *Trichoplusia ni* (insect) cell-lines (Ezure et al., 2006).

#### Chemical lysis

Chemical lysis relies on the use of enzymes or detergents to remove cell walls, typically used in the context of protein purification. Enzymes such as lysozyme (peptidoglycan layer in *E. coli)* or benzonase (nuclease to remove DNA and RNA) are commonly utilized in tandem with defined detergents such as Tween-20, Triton-X, or RIPA buffer or commercial mixtures such as BugBuster (Novagen) or CellLytic X (Sigma). To our knowledge, there is no successful production of a coupled cell-free system using chemical lysis, although attempts using lysozyme with freeze-thaw have been unsuccessful (Shrestha et al., 2012). While chemical lysis has been used for anaerobic cell-free activity assays (Kuchenreuther, Shiigi, & Swartz, 2014), no coupled transcription-translation has been demonstrated.

### Clarification

Following lysis, the resultant solution is typically extremely viscous and difficult to manipulate. For this reason, the lysis step is always followed by a clarification step in which the lysate is spun down in a centrifuge to separate cellular debris from the soluble substrates (active enzymes, small molecules, and co-factors, necessary to drive coupled transcription-translation). Although crude extract can be used with no clarification step, aside from issues arising from viscosity, background expression is increased relative to clarified extracts (T. W. Kim et al., 2006). Traditionally, clarification has consisted of (2x 30 min) 30,000 x g spins, a process that comprises a large portion of the processing time (resulting in term S30 extract) (Nirenberg & Matthaei, 1961). Two washes were later found to be unnecessary, with 1 wash sufficient to obtain equivalent signal (Liu et al., 2005).

In 2006 Kim (DM) and colleagues demonstrated a radical shift in clarification protocols by showing that one 12,000 x g spin for 10 minutes was successful in maintaining expression (T. W. Kim et al., 2006). Interestingly, cell-free expression from a 12,000 x g spin followed by no-dialysis was similar to that of the traditional 2 x 30,000 x g spins, and crude lysate with no processing showed only marginally less (20%) expression. This finding was reproduced independently, demonstrating a 30% increased yield using S12 over S30 (Pedersen, Hellberg, Enberg, & Karlsson, 2011). S12 preparations also demonstrated increased co-factors relative to S30 preparations (T.-W. Kim, Keum, et al., 2007a). S12 extract demonstrated workable viscosity and decreased background expression, but was strain specific to the Rosetta, BL21, and BL21-Star lines. Subsequently, 12,000 x g spins have become widely adopted for preparing cell-free systems from compatible strains (Kwon & Jewett, 2015; Shrestha et al., 2012; Sun et al., 2013).

### Runoff

A runoff reaction is typically conducted after clarification of the lysate, presumably to release ribosomes from bound mRNA and degrade leftover, sheared mRNA and DNA from the host strain (Jermutus, Ryabova, & Plückthun, 1998; Nirenberg, 1963). Before the runoff reaction, solutions are typically clear; afterwards, however, the solutions become cloudy, indicating degradation or modification (Sun et al., 2013). However, there has been little experimental evidence of this hypothesis, and it is a rich area of potential further exploration. Traditionally, the runoff reaction occurs at 37°C for 80 minutes, and mixes clarified lysate with a pre-incubation mix of Tris, Mg, ATP, DTT, amino acids, PEP, and pyruvate kinase. However, Swartz and colleagues first reported that the pre-incubation mix was unnecessary to obtain signal (Liu et al., 2005), and that a 37°C, 80-minute incubation of the post-clarified lysate was sufficient. In addition, ribosome release as the reason for the runoff is called into question, with a new hypothesis that the runoff activated activators or degraded inhibitors. Adding to the confusion, in 2015 Kwon and Jewett identified a strain-specific runoff property, with BL21-Star (DE3) strains not requiring runoff to activate protein production and other strains requiring different experimentally-optimized runoff steps (Kwon & Jewett, 2015). While current protocols for prototyping use a set 80-minute runoff without pre-incubation (Sun et al., 2013), this area is ripe for future research.

### Dialysis

After the runoff reaction, lysates are typically re-clarified by centrifugation to remove substrates accumulated during the reaction and dialyzed against a final S30 run buffer at 4 C (Sun et al., 2013). Traditionally, the dialysis step varies in length of time from one cycle of 3 hours (Sun et al., 2013) or 18 hours (Zubay, 1973) to 45 minutes x 4 cycles (Kigawa, Yabuki, Matsuda, Matsuda, Nakajima, Tanaka, & Yokoyama, 2004b). However, when explored with the runoff Swartz and colleagues found the dialysis step to be unnecessary, with no statistical difference between 0 - 4 dialysis cycles (Liu et al., 2005). This was confirmed by Kim (DM) and colleagues, who found dialysis unnecessary in the standard protocol, except when used after a 80 minute runoff step, presumably to remove by-products from the runoff (T. W. Kim et al., 2006). A potential added benefit of removing the dialysis step is the retention of cofactors that would otherwise pass through the 10kDa membrane used. There currently is a mix of protocols used, with some protocols utilizing dialysis (Garamella, Marshall, Rustad, & Noireaux, 2016; Sun et al., 2013; W. C. Yang et al., 2012) and others omitting dialysis (Kwon & Jewett, 2015; Shrestha et al., 2012). The effect of dialysis on extract composition and on prototyping ability is another area ripe for future research.

## E. coli ENERGY REGENERATION

The development of more efficient methods for energizing cell-free protein synthesis mirrors the maturation of cell-free extracts as a platform for synthetic biology (**Figure 3**). For decades, the ability to leverage the advantages of cell-free systems in industrial applications was limited by inefficient methods for regenerating ATP necessary for protein synthesis. In addition to being unable to sustain protein synthesis beyond an hour as a result of substrate instability (D. M. Kim & Choi, 1996; D.-M. Kim & Swartz, 2000b; R. G. Kim & Choi, 2000), early energy regeneration systems relied on prohibitively expensive substrates. These issues were gradually addressed by enabling and utilizing increasingly extended pieces of native cell metabolism to more efficiently drive protein synthesis. Much of the exploration in this area was not through building synthetic pathways, but rather through the observation that cell-free lysates innately conserve complex central metabolism, such as pathways for glycolysis and oxidative phosphorylation (**Figure 4**). For example, for oxidative phosphorylation to work, all enzymes in the TCA cycle and in the electron transport chain must be present and functional (Jewett et al., 2008; Jewett & Swartz, 2004a).

**Figure 3.**
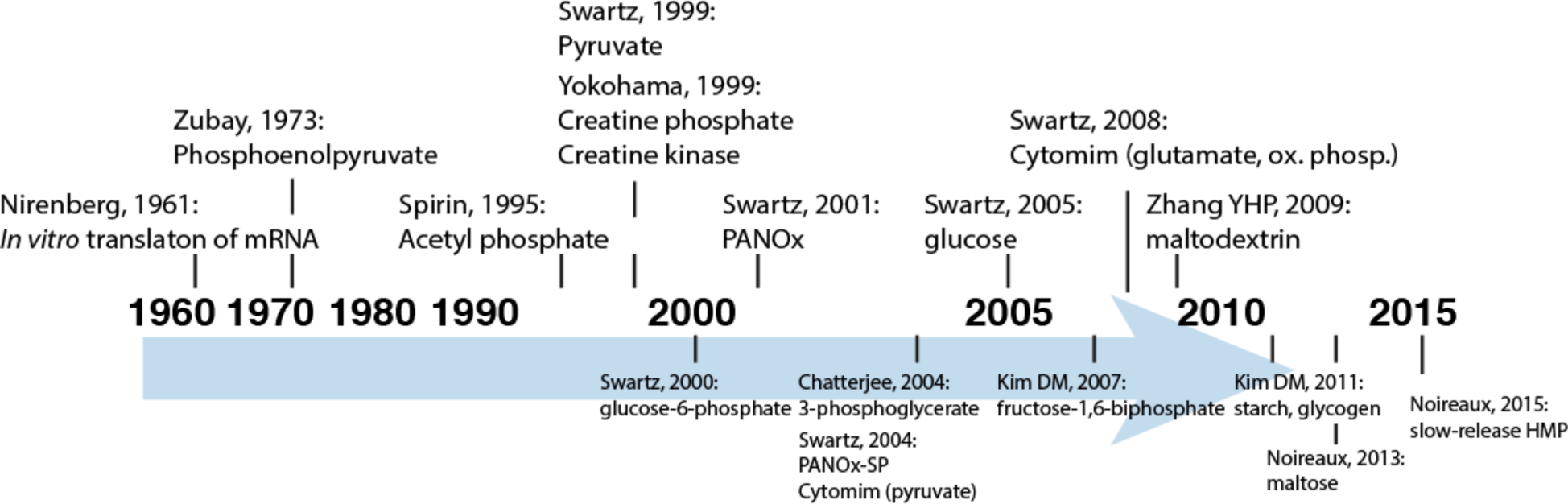
Energy sources to feed cell-free metabolism, arranged by year. Top of figure shows “major” breakthroughs in energy metabolism, while bottom of figure shows other breakthroughs.

**Figure 4.**
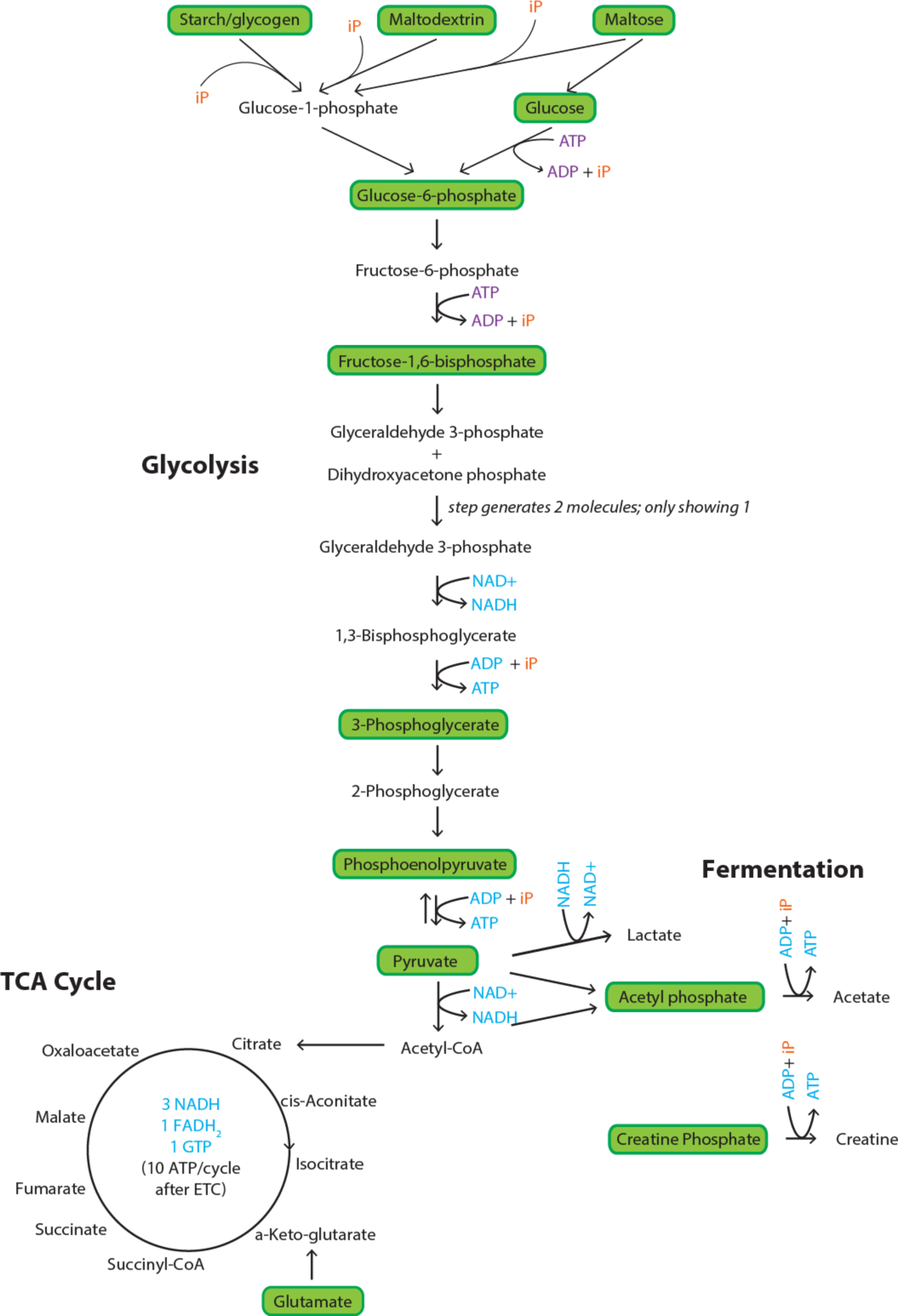
Simplified map of *E. coli* cell-free metabolism. Map is divided into Glycolysis, TCA Cycle, and Fermentation; areas in green are energy sources that have been explored for cell-free metabolism.

As a result, the latest cell-free systems feature a thousand-fold improvement in the relative cost of energy substrate demonstrated (Caschera & Noireaux, 2015) and protein synthesis can be extended to ten hours in simple batch mode (Caschera & Noireaux, 2014). This is a direct result of being able to exploit *E. coli* sugar metabolism in its entirety. The demonstration of such extensive, intact native machinery and the ability to manipulate its utility signals an important paradigm shift in cell-free systems. Rather than a black-box system used for simple protein production, cell-free extracts have evolved into a complex and valuable prototyping environment.

### General requirements

Although not constrained by energy costs associated with growth and maintenance in whole cells, cell-free extracts are still subject to stringent energy requirements posed by high-volume protein expression. Two molecules of ATP and two GTP are consumed in the formation of each peptide bond. Resource limitation is an important consideration in maximizing yields in protein production applications of cell-free reactions but is equally important in successfully implementing multi-step pathways and ensuring fidelity in rapid prototyping functions. Accordingly, a robust energy regeneration system is essential to maximizing extract performance across the board for all cell-free protein synthesis applications. This energy regeneration system must also be capable of avoiding inorganic phosphate accumulation (Spirin & Swartz, 2008) while maintaining pH within physiological range. We divide our discussion of energy regeneration in rough chronological order of development: single-step (substrate level) phosphorylation, multi-step pathway phosphorylation and oxidative phosphorylation.

### Single-step (substrate level) phosphorylation

The earliest iterations of cell-free extracts utilized molecules containing high-energy phosphate bonds as their source of energy (**Figure 5**). This paradigm remained relatively unchanged for many years. The most popular has been phosphoenolpyruvate (PEP) (Zubay, 1973). While PEP and pyruvate kinase (PK) together produced ATP, Spirin and colleagues hypothesized that acetyl phosphate could provide a cheaper alternative, and demonstrated equivalent signal with acetyl phosphate alone (Ryabova et al., 1995). This was the first evidence substrate-level phosphorylation was relatively independent of high-energy molecule chosen, and endogenous enzymes could be utilized.^1^ Four years later, Yokoyama and colleagues showed a completely exogenous system, creatine phosphate (CP), could also be used in conjunction with enzyme creatine kinase (CK) (Kigawa et al., 1999). CP/CK was tested after finding that PEP had inhibitory effects on cell-free reactions, which would later be attributed to inorganic phosphate accumulation from non-specific phosphatase degradation (D.-M. Kim & Swartz, 1999). Pyruvate kinase, creatine kinase, and acetate kinase each transfer their high-energy phosphate bonds to ADP to form ATP via substrate-level phosphorylation.

**Figure 5.**
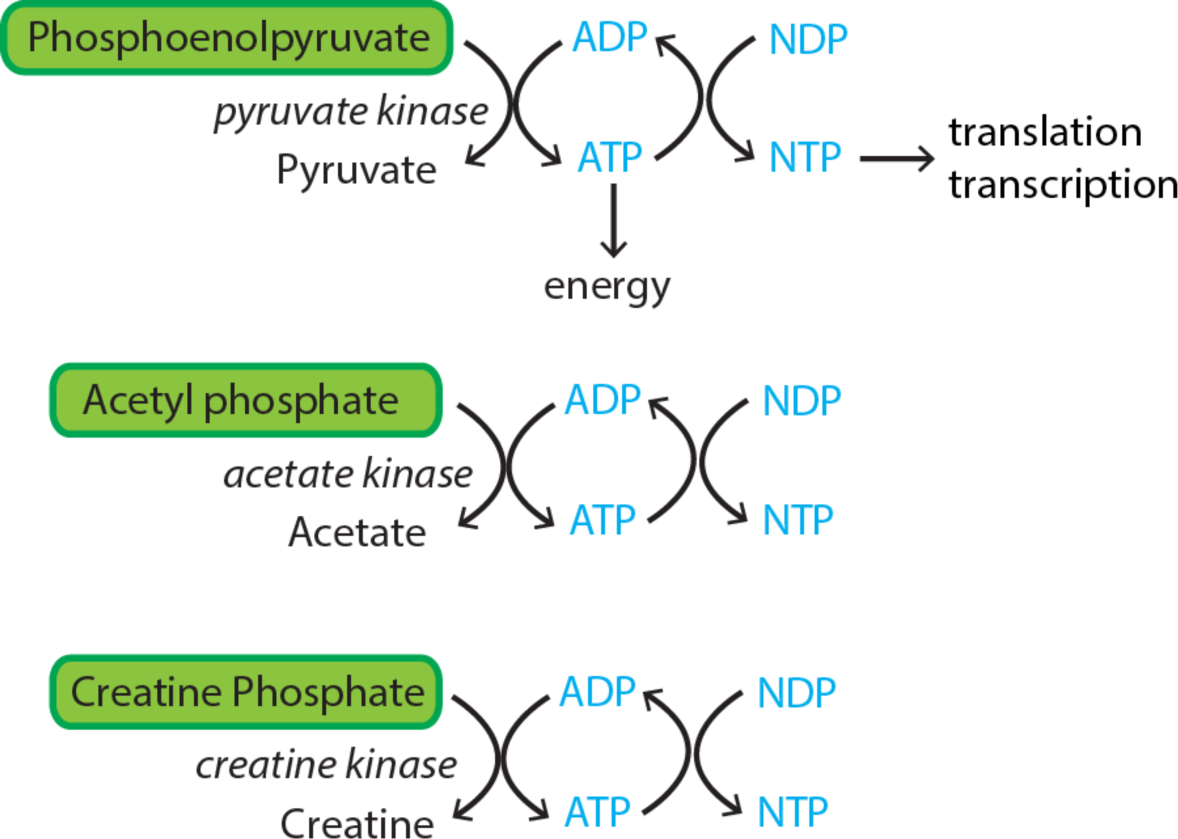
Substrate-level phosphorylation. Shown are three substrate-level phosphorylation modes utilized for cell-free systems: phosphoenolpyruvate/pyruvate kinase, acetyl phosphate/acetate kinase, and creatine phosphate/creatine kinase.

Although single-step phosphorylation of ATP is simple and effective in energizing cell-free protein synthesis, the use of PEP, CP, and AP has a number of drawbacks. The utility of high-energy phosphate molecules as energy donors is limited by their susceptibility to nonspecific attack by endogenous phosphatases (D.-M. Kim & Swartz, 1999; 2000b). The result is very transient expression as the energy molecules are quickly degraded. 70% of PEP was degraded into pyruvate and inorganic phosphate after a 30-minute incubation in S30 extract in the absence of DNA, indicating the presence of an unproductive sink for the supplied energy source (D.-M. Kim & Swartz, 1999). Protein yield is further limited by the accumulation of high concentrations of inorganic phosphate in solution resulting from the unproductive cleavage of the high-energy phosphate bonds. Reactions quickly terminate when phosphate concentrations reach 40-50 mM as a result of chelation of magnesium (T.-W. Kim, Oh, et al., 2007b), which is essential to biologically activating ATP and the function of essential enzymes (D.-M. Kim & Swartz, 1999). Altogether, this results in reactions not exceeding 1-2 hours in duration. To some extent, replenishing magnesium and the energy source in the reaction has been demonstrated to extend the duration of protein synthesis but such an approach rules out simple batch-mode reactions (D.-M. Kim & Swartz, 2000b). Another solution explored addition of inorganic phosphate and glucose to the growth medium in which the cells are grown, which limited phosphatase activity in extracts by suppressing expression of phosphatases during growth (R. G. Kim & Choi, 2000).

### Multi-step pathway phosphorylation

A number of different systems from 1999 onwards were developed to address the weaknesses inherent in systems dependent on high-energy phosphate-bonded compounds. These strategies relied on utilizing multi-step enzymatic pathways in order to more efficiently harness the energy of the high-energy phosphate compounds. By utilizing multi-step pathways, substrates would be less prone to phosphatase attack and a spike in inorganic phosphate concentrations. In addition, ATP generation could be extended over the course of the reaction. The first multi-step system generated acetyl phosphate through the addition of pyruvate and pyruvate oxidase in the presence of thiamine pyrophosphate (TPP) and flavin adenine dinucleotide (FAD) (D.-M. Kim & Swartz, 1999). Substrate-level phosphorylation of acetyl phosphate produces ATP by endogenous acetate kinase. This method is more resistant to phosphatase activity as a lower, sustained concentration of acetyl phosphate is lower relative to the K_m_ of the phosphatases. Importantly, the use of an un-phosphorylated substrate serves as an inorganic phosphate sink, thereby preventing phosphate buildup by coupling the activity of pyruvate oxidase to the inorganic phosphate produced by acetyl kinase.

In 2001, the PANOx system (PEP, amino acids, NAD+, oxalic acid) debuted as a highly rationally-engineered multi-step pathway phosphorylation system (D.-M. Kim & Swartz, 2001) (**Figure 6**). PANOx employs multiple enzymatic steps to more efficiently harness the energy of high-energy phosphorylated molecules. This system utilizes PEP as the main source of energy but benefits from substrate-level phosphorylation at two points: the conversation of PEP to pyruvate, and the conversion of acetyl phosphate to acetate. The latter conversion is enabled by the addition of NAD+ and CoA, which drive the conversation of pyruvate to acetate. Addition of oxalic acid prevents the nonproductive reverse reaction of pyruvate to PEP by inhibiting the activity of PEP synthase (D.-M. Kim & Swartz, 2000a), thereby driving flux forward. Finally, amino acids were supplemented to replace degradation of arginine, serine, tryptophan, and cysteine (Michel-Reydellet et al., 2004). The second generation of this system, PANOx-SP adapted the PANOx environment to look more like the cell by replacing polyethylene glycol with spermidine and putracine (the SP of PANOx-SP), and removing HEPES buffer (Jewett & Swartz, 2004a). These changes were made to encourage metabolism of pyruvate, as pyruvate alone provided only 20% of the signal of PEP. The PANOx, PANOx-SP, and variants thereof are still widely in use as energy regeneration methods, both in academic settings (T Michaele Holland & Bundy, 2012; Hong et al., 2014; Kwon & Jewett, 2015) and in commercial settings (Roche, now Biotechrabbit RTS-100).

**Figure 6.**
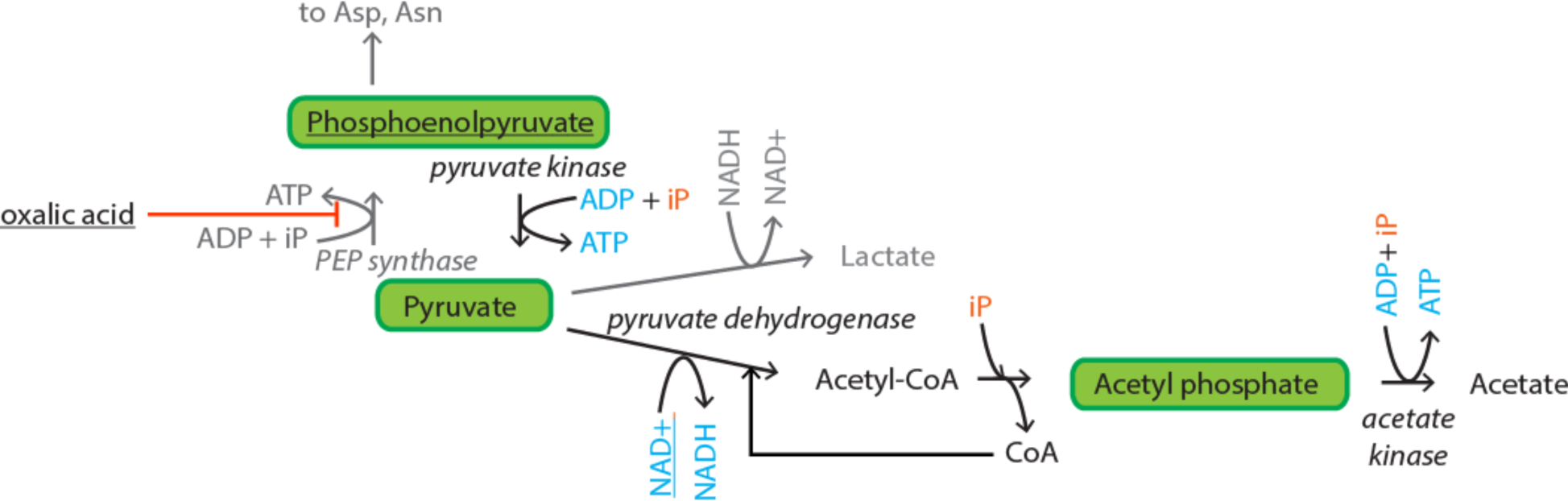
PANOx energy regeneration system. Underlined are PANOx additives (amino acids not shown). In grey are non-productive pathways. In italics are enzymes. ATP is generated from conversion of PEP to pyruvate via pyruvate kinase, and converstion of acetyl phosphate to acetate via acetate kinase. Figure adapted from (Jim Swartz, 2006)

Also in 2001, glucose-6-phosphate (G6P) was found as sufficient to energize cell-free reactions (D.-M. Kim & Swartz, 2001). This finding was particularly notable, as G6P is 9 steps removed from pyruvate in the glycolytic pathway. At worst case, G6P required conversion to fructose-1,6-biphosphate (F1,6-BP) before substrate-level phosphorylation; at best case, the catabolic machinery for glycolysis remained intact in cell-free extracts. Logically, it follows that any of the intermediates in the glycolytic pathway can be utilized as the starting substrate for ATP regeneration. Following this line of reasoning, 3-phosphoglycerate (3-PGA), a glycolytic intermediate two steps upstream of PEP, was employed as a primary energy source to offer further improvement on the existing ATP regeneration paradigm. By co-opting endogenous enzymes to generate PEP *in situ* from 3-PGA, Chatterjee and colleagues were able to extend the duration of protein synthesis well beyond the limit of an analogous system utilizing PEP (Sitaraman et al., 2004). The continuous synthesis of PEP allowed the system to evade the phosphatase activity that hampered ATP regeneration by maintaining PEP at low enough concentrations to avoid premature degradation. The 3-PGA system would form the basis of prototyping extracts from Noireaux, Murray, and colleagues (Shin & Noireaux, 2010; Sun et al., 2013). Similarly, F1,6-BP, which is further upstream than 3-PGA, was demonstrated to outstrip 3-PGA as an energy donor as theoretically one molecule of FBP would yield more ATP than intermediates downstream (T.-W. Kim, Keum, et al., 2007a).

The glycolytic pathway, in its entirety, has also been leveraged for the cheap and efficient replenishment of ATP. In such a situation, glucose, the cost of which is negligible relative to most other energy sources, can serve as an economical and efficient substrate for energizing protein expression. In contrast to traditional substrate-level phosphorylation reactions that yield only one mole of ATP per mole of costly energy molecule, one mole of glucose can yield 2 or 3 moles of ATP, respectively. A landmark moment was the usage of glucose, notable as a cheap substrate, for energizing cell-free systems in parallel with NMPs (vs. NTPs) (Calhoun & Swartz, 2005c). While effective, glucose metabolism was found to decrease the pH of the reaction below the physiological range as a result of organic acid production (Calhoun & Swartz, 2005b), thereby requiring extensive buffering. Reactions utilizing glucose required the addition of inorganic phosphate in addition to pH buffering to express on par with G6P. One approach coupled glucose metabolism with creatine phosphate and creatine kinase in a complementary energy regeneration system (T.-W. Kim, Oh, et al., 2007b). The normally inhibitory inorganic phosphate from creatine phosphate metabolism served as the phosphate source necessary to activate glucose metabolism. Although the amount of protein yielded by this combination of resources was high, the return to creatine phosphate offset this advantage with a much higher cost to yield ratio. Another iteration refined the parameters for glucose utilization and dramatically extended reaction duration and productivity, resulting in a six-hour reaction yielding 1.8 mg/mL of protein (T.-W. Kim, Kim, Oh, & Kim, 2008). This was accomplished by growing the cells in the presence of glucose and phosphate, further fortifying the pH buffering capacity, and utilizing the S12 extract preparation method to preserve cofactors.

Recently, cell-free systems have been engineered to use complex sugars in order to prevent pH issues, maintain cost advantage, and allow for long-lasting energy release. Wang and Zhang (YHP) in 2009 demonstrated that maltodextrin, in combination with supplemental maltodextrin phosphorylase and phosphoglucomutase, could effectively energize cell-free reactions at very low cost (Y. Wang & Zhang, 2009). This strategy integrates phosphorolysis to serve as an inorganic phosphate sink and glycolysis in addition to the PANOx pathway to generate ATP. Each glucose equivalent in maltodextrin can produce one more net ATP relative to glucose by consuming inorganic phosphate rather than ATP in the formation of G6P. Maltodextrin resulted in less pH perturbations (relative to glucose, PEP, and G6P in analogous systems), and in more homeostatic and stable reaction conditions. Further refinement of the polysaccharide approach utilizing starch and glycogen demonstrated that protein synthesis could continue for 12 hours in a simple batch mode, yielding 1.7 mg/mL of protein without addition of exogenous enzymes (H.-C. Kim, Kim, & Kim, 2011). Kim et al. also demonstrated the maintenance of a steady supply of ATP without drastic alteration of pH which they postulated might explain the improved solubility of synthesized protein. Quantification of the ATP and starch levels after cessation of transcription at 12 hours showed that only 20% of the starch had been consumed and ATP concentrations were still constant, implying that ATP supply was not the limiting factor. In an effort to improve upon the system by introducing a gradually released phosphate reservoir, bypassing the presence of potentially inhibitory amounts of inorganic phosphate, hexametaphosphate was recently utilized in place of potassium phosphate (Caschera & Noireaux, 2015). An approach combining the strengths of the 3-PGA system with maltose, a disaccharide acting as an inorganic phosphate sink and secondary energy source was also pursued (Caschera & Noireaux, 2013).

### Oxidative Phosphorylation

The Cytomim system was originally produced to metabolize (cheaper) pyruvate in lieu of (more expensive) PEP, with the working hypothesis that conditions more representative of the cytoplasm would be necessary for pyruvate utilization (Jewett & Swartz, 2004b; 2004a). Interestingly, however, Jewett and Swartz discovered that ATP generation and protein synthesis continued beyond the depletion of pyruvate, the presumed energy substrate. On follow-up, the depletion of glutamate and formation of TCA cycle intermediates was observed, demonstrating that glutamate alone could serve as a stand-alone energy substrate (Jewett et al., 2008). This process was found to be heavily oxygen-dependent, thereby confirming that oxidative phosphorylation could be activated in cell-free extracts. Biochemical inhibitors of the electron transport chain also significantly reduced the protein yielded by the cell-free system. This represents a substantial shift in thinking, as the TCA cycle as well as the electron transport chain are necessary. It was theorized that lysis methods causing high shear rates (eg. high-pressure homogenization) allows inverted membrane vesicles (IMVs) upon which oxidative phosphorylation can occur.

### Energy Regeneration in the context of Synthetic Biology

While there has been extensive innovation in energy regeneration in cell-free systems, less clear are the conditions that are necessary to enable synthetic biology applications to function, such as the rapid prototyping of circuits (Garamella et al., 2016; Sun et al., 2014; Takahashi et al., 2015) and of pathways (Karim & Jewett, 2016; Wu, Culler, Khandurina, Van Dien, & Murray, 2015). For circuit prototyping specifically, one can assume that interactions such as protein-binding strength to operators, or weak K_m_ binding events, are more critical than pure protein expression. In addition, native polymerases are favored over T7 polymerase to better emulate cellular conditions (Shin & Noireaux, 2010). With a goal of prototyping to match cellular function and implement complexity (versus pure protein production), it is likely that re-evaluation of existing approaches will be necessary to support this new application. To date, two protocols have been used for circuit prototyping: (1) a protocol utilizing bead-beating and 3-PGA energy regeneration (Sun et al., 2013), and (2) a protocol mixing bead-beating or French-press preparation and 3-PGA with maltodexrin and/or maltose (Garamella et al., 2016). In addition, the protocol of (Sun et al., 2013) has been used for prototyping pathways for 1,4-BDO (Wu et al., 2015) and violacein (Nguyen, Wu, Guo, & Murray, 2015), as well as a modified PANOx-SP run off of T7 RNA polymerase for n-butanol (Karim & Jewett, 2016). However, there has been no published work on engineering of the cell-free protocol to specifically support circuit prototyping.

While we now know that cell-free systems contain large amounts of intact metabolism, there is also a need to apply ‐omics technologies (genomics, proteomics, metabolomics, transcriptomics, glycomics) to better understand the cell-free “black-box.” Since the dense period of discovery from 1999 - 2011, there has been a lack of published work seeking to understand the extent to which central metabolism can be activated and manipulated. This is particularly compelling with the new tools available since the bulk of discovery was conducted, including RNAseq (Mortazavi, Williams, McCue, Schaeffer, & Wold, 2008), high-throughput gene synthesis and assembly (Kosuri et al., 2010), and high-throughput gene sequencing (Shendure, Mitra, Varma, & Church, 2004). It is also increasingly evident that cell-free systems are not uniform, standard “collections” of lysates, but rather complex compositions that are affected by the multiple variables of preparation and energizing. These complex compositions may require standardization of preparation, or individual analysis per batch to understand variability that result extract-to-extract (Takahashi et al., 2014; 2015). Attempts by Panke and colleagues to conduct real-time analysis on lysates is a start at understanding this complexity (Bujara, Schümperli, Pellaux, Heinemann, & Panke, 2011).

## ACKNOLEDGEMENTS

The authors declare a conflict of interest: ACC, RMM, ZSS hold ownership in Synvitrobio, Inc. The work presented here was funded off of a DARPA SBIR to Synvitrobio, Inc. (ACC, ZSS), contract No: W911NF-16-P-0003, and a Caltech Grubstake Grant (ACC, RMM, ZSS). The views and conclusions contained in this document are those of the authors and should not be interpreted as representing officially policies, either expressly or implied, of the Defense Advanced Research Projects Agency or the U.S. Government.

## Footnotes

1 It is likely that if PEP was supplied alone, the system would still produce ATP from the endogenous pyruvate kinase present in the extracts.

